# Visualizing tumor evolution with the fishplot package for R

**DOI:** 10.1101/059055

**Authors:** Christopher A. Miller, Joshua McMichael, Ha X. Dang, Christopher A. Maher, Li Ding, Timothy J. Ley, Elaine R. Mardis, Richard K. Wilson

## Abstract

**Background:** Massively-parallel sequencing at depth is now enabling tumor heterogeneity and evolution to be characterized in unprecedented detail. Tracking these changes in clonal architecture often provides insight into therapeutic response and resistance. Easily interpretable data visualizations can greatly aid these studies, especially in cases with multiple timepoints. Current data visualization methods are typically manual and laborious, and often only approximate subclonal fractions.

**Results:** We have developed an R package that accurately and intuitively displays changes in clonal structure over time. It requires simple input data and produces illustrative and easy-to-interpret graphs suitable for diagnosis, presentation, and publication.

**Conclusions:** The simplicity, power, and flexibility of this tool make it valuable for visualizing tumor evolution, and it has potential utility in both research and clinical settings. Fishplot is available at https://github.com/chrisamiller/fishplot

## Background

Most cancers are heterogeneous and contain multiple subclonal populations that can be detected via high depth massively parallel sequencing. An increasing number of studies are collecting and sequencing longitudinal samples, allowing the clonal evolution of tumors to be tracked in detail. Though tools have been developed for inferring subclonal architecture[1–4] and for determining tumor phylogeny[5–7], few offer compelling and intuitive visualizations.

We reported one of the first studies describing tumor evolution defined by whole genome sequencing, in patients with relapsed Acute Myeloid Leukemia, and that publication contained a series of custom figures showing changes in clonal architecture between the primary and relapse presentations[8]. Often called “fish plots” due to their resemblance to tropical fish, these visualizations have become widely adopted, both by our group and others[9–12]. Until now, each has been created in vector-art programs like Adobe Illustrator, which is laborious and makes representing accurate proportions challenging. As the sizes of cohorts have grown, this approach has quickly become untenable.

To enable the creation of these plots in a robust and automatable fashion, we have developed an R package (“fishplot”) that takes estimates of subclonal prevalence at different timepoints, and outputs publication-ready images that accurately represent subclonal relationships and their relative proportions. Fishplot is available at https://github.com/chrisamiller/fishplot.

## Implementation

The fishplot package was implemented in R, and requires a minimal set of dependencies (the “plotrix”, “png”, and “Hmisc” packages). Several inputs, including the clonal fractions of each tumor cell population, a representation of descent in the form of parental relationships, and the timepoints at which the samples were obtained are required. These data are readily available from existing tools, such as the clonevol package, which already includes code that interfaces with fishplot for visualization[7]. This feature enables seamless integration into existing genomic analysis pipelines. Figures are output through the R standard graphics libraries, which allow for the creation of vector or raster-based images of any size, suitable for a wide range of applications.

## Results/Discussion

We have applied fishplot to a number of different cancer genomics studies, and three representative results are displayed in Figure 1.

**Figure 1:**
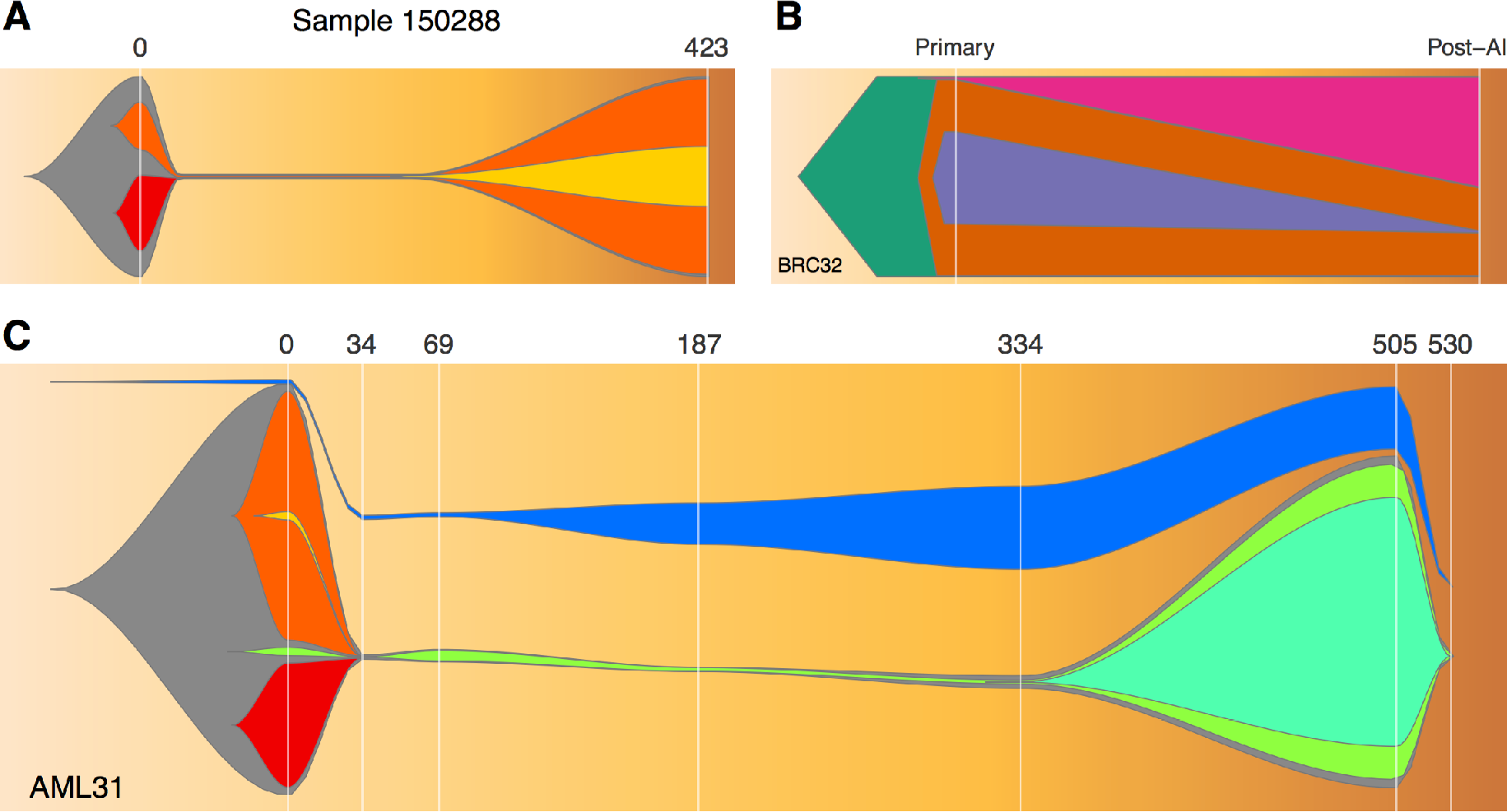
Visualizing tumor evolution with the fishplot package. Panel A: A case of primary and relapsed AML. Panel B: A breast cancer before and after neoadjuvant aromatase inhibitor therapy. Panel C: An AML with complex clonal structure and 7 timepoints.

We first created a fish plot of an Acute Myeloid Leukemia patient with a chemotherapy-induced bottleneck that eliminated one subclone, while another survived and drove the relapse (Figure 1A). This plot uses the default color scheme and the default curve splining method for smoothing, along with timepoint labels representing the number of days since tumor presentation.

We next plotted a breast tumor sequenced before and after 4 months of neoadjuvant aromatase inhibitor therapy [Miller, et al in press] (Figure 1B). This therapy did not induce an extreme bottleneck, but nonetheless resulted in substantial clonal remodeling. The resulting figure uses user-defined colors and represents subclones as polygons, without curve smoothing. The standard numeric labels were replaced with categorical labels, but the timepoints remain scaled appropriately.

Lastly, we created a model of AML31, a patient that was sampled with ultra-deep sequencing at many timepoints, allowing even very rare (<1% VAF) subclones to be detected[10] (Figure 1C). Chemotherapy did not completely eliminate the cancer in this case, resulting in detectable levels of tumor until relapse at day 505 (and subsequent clearance of the tumor with salvage chemotherapy). The fishplot shows this progression, including the failure to completely clear the tumor. This patient also had oligoclonal skewing post-chemotherapy, resulting in a non-cancerous hematopoetic stem cell expansion of a specific TP53-containing cell that was not genetically related to the patient’s tumor (Figure 1C, blue). The package includes functionality for representing such unrelated clones, which is also useful in the case of “collision tumors” with independent origins. The code used to produce Figure 1 is available as supplemental data and can be also found within the example scripts in http://github.com/chrisamiller/fishplot/tests/tests.R

## Conclusions

Characterizing subclonal architecture and the ways in which tumors evolve, both over time and in the context of therapeutic intervention, is important for understanding therapy resistance, which contributes to tens of thousands of cancer deaths each year. Fish plots, like those presented here, provide researchers and clinicians with an intuitive and accurate understanding of how an individual tumor is changing over time, making analysis and diagnosis easier. Despite being designed for tracking tumor evolution, this tool may also find niches outside of cancer biology, and could easily be used to represent the changing landscapes of microbial populations, for example. Our group has already used images created by the fishplot package in a number of genomic pipelines and pending publications, and we anticipate that it will be adopted widely within the large community of scientists studying tumor evolution.

## Acknowledgements

Research reported in this publication was supported in part by funding provided by grant U54HG003079 from the National Human Genome Research Institute to RKW.

